# Absence of a faster-X effect in beetles (*Tribolium*, Coleoptera)

**DOI:** 10.1101/754903

**Authors:** Carrie A. Whittle, Arpita Kulkarni, Cassandra G. Extavour

**Affiliations:** Department of Organismic and Evolutionary Biology, Harvard University, 16 Divinity Avenue, Cambridge MA 02138, USA; Department of Molecular and Cellular Biology, Harvard University, 16 Divinity Avenue, Cambridge MA 02138, USA

**Author notes:** Corresponding Author: Cassandra G. Extavour.

**Keywords:** *Tribolium castaneum*, faster-X, sex-biased expression, dosage compensation, dN/dS

## Abstract

**Background:** The faster-X effect, namely the rapid evolution of protein-coding genes on the X-chromosome, has been reported in numerous metazoans. However, the prevalence of this phenomenon across metazoans and its potential causes remain largely unresolved. Analysis of sex-biased genes may elucidate its possible mechanisms: a more pronounced faster-X effect in male-biased genes than in female-biased or unbiased genes, suggests fixation of recessive beneficial mutations rather than genetic drift. Further, theory predicts that the faster-X effect should be promoted by X-chromosome dosage compensation, but this topic remains rarely empirically examined.

**Results:** Here, we asked whether we could detect a faster-X effect in genes of the beetle *Tribolium castaneum* (and *T. freemani* orthologs), which has X/Y sex-determination and heterogametic males. Our comparison of protein sequence divergence (dN/dS) on the X-chromosome versus autosomes indicated the complete absence of a faster-X effect. Further, analyses of sex-biased gene expression revealed that the X-chromosome was strongly enriched for ovary-biased genes, which evolved under exceptionally high constraint. An evaluation of male X-chromosome dosage compensation in the gonads and in non-gonadal somatic tissues showed an extreme lack of compensation in the testis. This under-expression of the X chromosome in males may limit the phenotypic effect, and therefore likelihood of fixation, of recessive beneficial X-linked mutations in genes transcribed in male gonads.

**Conclusions:** We show that these beetles display a rare unequivocal example of the absence of a faster-X effect in a metazoan. We propose two potential causes for this, namely high constraint on X-linked ovary-biased genes, and an extreme lack of dosage compensation of genes transcribed in the testis.

## Background

The “faster-X” effect, that is, the rapid evolution of protein-coding genes on the X chromosome, has been widely reported in a range of metazoan systems with sex chromosomes [1, 2]. Higher rates of protein divergence of genes on the hemizygous X-chromosome (faster-X, or faster-Z in W/Z systems) than on autosomes has been observed in organisms including primates [3, 4], humans [4], rodents [5], birds [6, 7], moths [8], aphids [9], and very recently in spiders [10]. In other organisms, however, a faster-X effect is more ambiguous. For example, signals of this effect have sometimes, but not always, been observed in studies of fruit flies [2, 11, 12], and variable results on the presence or strength of the faster-X effect have been reported in butterflies [13, 14].

With regards to the mechanisms that might account for the faster-X effect, it has been proposed that X-linked genes might evolve faster in protein sequence than those on autosomes due to efficient fixation of recessive beneficial mutations in the hemizygous state, a notion that has found empirical support in some animal taxa [1, 2, 4, 5, 11, 15]. An alternative mechanism is that the effect results largely from fixation of recessive, mildly deleterious mutations via genetic drift. Studies of birds and aphids support this mechanism, which has been suggested to be facilitated by the lowered effective population size of the X chromosome [1, 7, 9, 16].

The study of sex-biased gene expression, that is, those genes preferentially upregulated in one sex, has helped to decipher the forces shaping the molecular evolutionary rates on the X-chromosome versus autosomes [17–20], and thus to better understand the faster-X effect [7, 8, 12–16]. For instance, under a model wherein the faster-X effect is caused by rapid fixation of beneficial mutations in the hemizygous state, in organisms where males are the heterogametic sex, this effect is predicted to be strongest in male-biased genes, and relatively lower in female-biased and unbiased genes [16]. This prediction is based on the hypothesis that X-linked recessive beneficial mutations should largely exert their fitness effects in males, as their hemizygous state would preclude the possibility of non-mutant alleles masking the phenotypic effect of such recessive mutations [7, 15, 16]. Empirical support from this model comes from a study of *Drosophila*, in which assessment of protein divergence (dN/dS) of genes on the X-chromosome and autosomes, revealed a faster-X effect for all three classes of sex biased genes (male-biased, female-based and unbiased). In this study, protein sequence divergence was highest in male-biased genes and lowest in female-biased genes, empirically supporting a model of fixation of beneficial mutations on the X chromosome [15, 16]. In chickens, which have WZ sex chromosomes and female heterogamy, elevated dN/dS has been reported across all studied genes on the Z-chromosome, consistent with the faster-X (or faster-Z in this case) effect [21]. However, the prediction of higher dN/dS for female-biased genes on the Z-chromosome was not met in this study, suggesting that the faster-Z in these birds was not due to fixation of recessive beneficial mutations, and rather might be attributable to fixation of neutral or slightly deleterious mutations via genetic drift [7, 16]. Recently, similar results were reported for the W/Z chromosomes of *Heliconius* butterflies [14]. At present however, the study of the faster-X effect, including the role of sex-biased gene expression, remains limited to just a few model organisms, and the putative underlying mechanisms appear to be variable.

The faster-X effect may be expected to be most strongly observed in organisms with complete dosage compensation, wherein expression levels of X-linked genes are upregulated in the heterogametic sex, such that the X to autosome ratio (X:A) is one or close to one [1, 2, 22, 23]. Under this hypothesis, in organisms with incomplete X-chromosome dosage compensation, such that X:A <1, X-linked recessive beneficial mutations would have relatively low expression levels, and thus putatively weak phenotypic effects (or selection coefficients[1]), in the hemizygous sex. This could make beneficial mutations exposed on the single male X-chromosome unlikely to be fixed any more frequently than they would be if they were autosomal, possibly resulting in evolutionary rates of X-linked genes that are not significantly faster than those of autosomal genes [1, 7]. Support for the notion that dosage compensation may impact the rates of X-(or Z-) linked gene evolution is provided by the observation that certain butterflies display incomplete dosage compensation [24]; this correlates with, and thus may contribute towards, the observed lack of an elevated faster-Z effect in female-biased genes in those organisms [14, 24]. Similar patterns have been described in birds [7]. At present however, the relationship between faster-X effect and dosage compensation remains rarely empirically evaluated.

An important factor to consider in the study of dosage compensation, is that this phenomenon may vary among tissues within an organism. For instance, studies in *Drosophila* have shown that complete dosage compensation of X-linked genes is observed in male somatic tissues, but not in the testis [25–27]. In the context of these findings in *Drosophila*, it appears possible that the degree of gonadal and somatic dosage compensation could in theory [1] influence its observed faster-X effect [2, 11, 12]. Thus, given that dosage compensation may vary among tissues, and particularly in the gonads, specific study is warranted of the faster-X effect and its association to gonadal and non-gonadal dosage compensation in animal models.

A model insect genus that offers new opportunities to study the faster-X effect is the beetle system *Tribolium* (Coleoptera). Coleoptera is the largest insect order, with recent estimates of over 1.5 million species, thus comprising approximately 40% of all arthropod species [28]. The rust red flour beetle *T. castaneum* is a well-established model system for genetics and for the evolution of developmental mechanisms [29–33], and has extensive genomic resources available for research [34–36]. In addition, its less well-studied sister species *T. freemani*, from which it diverged approximately 12-47 Mya, comprises a suitable system for comparative genomic study [37]. To date, however, to our knowledge the primary genome-wide sex-biased expression research in *Tribolium* that includes X-chromosome analyses consists of a foundational study based on whole male versus whole female contrasts and microarray data in *T. castaneum* [38]. That assessment made several significant findings, including that the female-biased genes were highly overrepresented on the X-chromosome [38], which was proposed to be explained by a mechanism of overexpression of X-linked genes in females as an imperfect response to male dosage compensation [38]. In addition, the study’s authors reported that X-linked genes with male-biased expression were comparatively uncommon, a trend also observed in other organisms such as *Drosophila* [38]. In addition to that assessment, other transcriptome studies in *Tribolium* include a recent study using RNA-seq in *T. castaneum* that examined differential expression among somatic, germ line, and embryonic tissues [39]. The study reported identification of potentially useful transcripts and genes for generating genetic constructs for the investigation of development and pest control in this species [39]. A separate investigation of codon and amino acid usage was also conducted across the *T. castaneum* genome with respect to gene expression [35]. None of these studies, however, assessed evidence for or against the faster-X effect in *Tribolium*. Moreover, to our knowledge, there have been no between-species analyses of protein sequence divergence (dN/dS), and its potential relationship to sex-biased gene expression and dosage compensation. Thus, we considered that original analyses addressing these important topics in this beetle model could provide valuable insights into the breadth of the faster-X phenomenon across animals, and help decipher its underlying mechanisms.

Here, we describe a rigorous assessment of the faster-X effect in *T. castaneum*, including evaluation of its relationship to sex-biased gene expression and dosage compensation, using newly generated RNA-seq data from gonads and gonadectomized (GT-) males and females. Our assessment of dN/dS in 7,751 *T. castaneum* genes with high confidence orthologs in its sister taxon *T. freemani* reveals the complete absence of a faster-X effect in this taxon. Instead, we find a slower rate of evolution of X-linked as compared to autosomal genes. Further, we show that the faster-X effect is not found at any level for male-biased, female-biased or unbiased genes identified from the gonads and from non-gonadal somatic tissues. We demonstrate that the slow-X effect in this taxon is largely due to the slow sequence evolution of ovary-biased genes located on the X-chromosome, which are more common, and are more highly constrained, than those on autosomes. Moreover, with respect to dosage compensation, we report that GT-males exhibit high X-chromosome dosage compensation with an X/A ratio near one. However, an extreme absence of dosage compensation is evident for hemizygous X-linked genes expressed in the testis. We suggest that this may give rise to weak phenotypic effects of such genes [1], potentially limiting fixation of recessive beneficial mutations when transcribed in male gonads, thereby impeding a faster-X effect. Our results thus provide additional empirical support [7, 14] for a notion that has previously been proposed theoretically [1, 2, 7, 22]. Taken together, we propose that the unusual absence of a faster-X effect in these beetles may be influenced by two major phenomena: (1) the accumulation of highly constrained ovary-biased genes on the X-chromosome, and (2) the lack of dosage compensation in the male gonads, which may act to minimize fixation of recessive beneficial mutations of genes transcribed in these tissues.

## Results

The complete list of previously annotated [36] protein-coding genes in our main target taxon *T. castaneum* were downloaded for study (N=16,434 genes, Ensembl Metazoa (http://metazoa.ensembl.org). Using the full CDS list (longest CDS per gene) we identified 7,751 high confidence orthologs in its sister species *T. freemani* for our study of protein sequence evolution (dN/dS, see Methods; note that expression results for all 16,434 *T. castaneum* genes are described throughout when appropriate). The use of closely related sister species is a common approach to study the protein sequence divergence of sex-biased genes in metazoan models (*cf.* [6, 9, 15, 40–42]). Values of dN/dS <1, =1, and >1 suggest that purifying, neutral and positive selection respectively are likely to predominantly shape the evolution of protein coding genes [43]. However, even when dN/dS <1 (as is typical in gene-wide analysis), relatively elevated values suggest reduced constraint, which could be due to relaxed selection and/or adaptive evolution. Under the faster-X effect, dN/dS is predicted to be elevated for protein-coding genes on the X-chromosome as compared to those on autosomes [7, 15, 16, 21].

We first assessed whether this beetle system exhibited a faster-X effect. Box plots of dN/dS of genes located on the X-chromosome versus autosomes are shown in Fig. 1, showing no tendency for higher dN/dS in genes on the X chromosome. In fact, we observed the opposite: dN/dS was statistically significantly lower for X-linked genes than for autosomal genes in this taxon (MWU-test P=0.002). From a total of 432 studied X-linked genes and 7,319 autosomal genes distributed across nine autosomes, the median dN/dS values were 0.686 and 0.906 respectively (Fig. 1A), yielding a ratio of dN/dS values for the X-chromosome to autosomes across all genes (X/A_dN/dS (all genes)_) of 0.76 (Fig. 1B). Thus, the X/A_dN/dS (all genes)_ value is considerably below 1, a result opposite to the >1 value expected under a faster-X effect [4, 14-16, 21]. Further, the mean dN/dS on the X-chromosome was about half (ratio of 0.54) that observed on autosomes (Fig. 1B). Thus, these results indicate the absence of a faster-X effect in this taxon, differing from that observed in most other metazoan systems studied to date. Together, these data show a slower-X pattern in *Tribolium*.

**Fig. 1.**
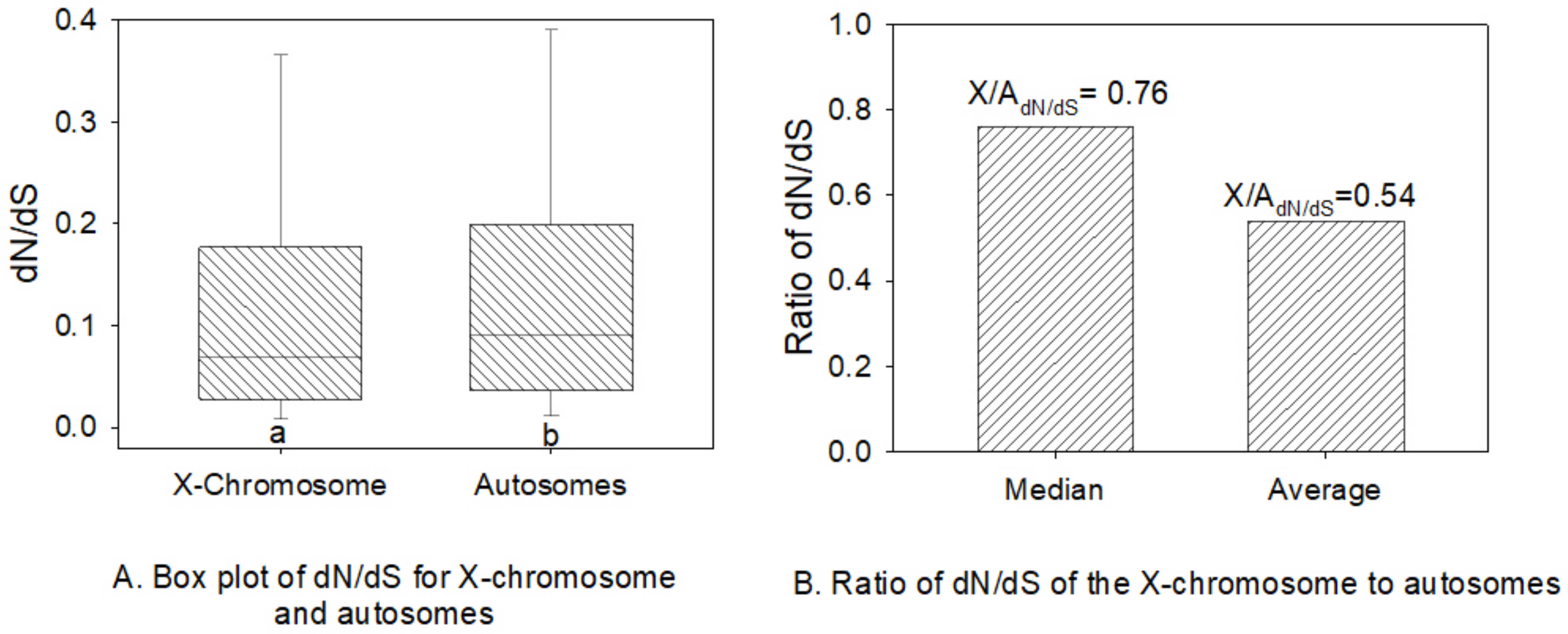
The dN/dS of genes located on the X-chromosome versus autosomes. A) Box plots of dN/dS showing the median, upper and lower quartiles, and 95/5th percentiles; B) the ratio of dN/dS for the X-chromosome versus the autosomes using the median and mean values per group. Different letters under bars in panel A indicate a statistically significant difference using MWU-tests.

### Assessment of sex-biased genes on the X-chromosome versus autosomes

Sex-biased gene expression has primarily been used to help discern the potential causes of the faster-X (or -Z) after it has been detected in an organism [7, 12–16]. Having observed no evidence of a faster-X effect for this beetle taxon, we asked if sex-differences in gene expression could help suggest mechanisms that might explain the absence of this effect (Fig. 1). We also wished to determine whether a faster-X effect was present for specific categories of sex-biased genes, including male-biased, female-biased and unbiased genes. For this assessment, we generated new large-scale RNA-seq datasets for adult male testes and ovaries, and for the gonadectomized bodies of adult males and females (hereafter referred to as GT-males and GT-females respectively, or non-gonadal somatic tissues) (Table S1). We mapped reads to annotated *T. castaneum* genes (See Methods), and identified sex-biased genes for the gonads (testis versus ovary) and for the GT-soma (GT-males versus GT-females) as those with a two-fold and statistically significant difference (P<0.05) in expression using Deseq2 [44]. We found that 25.8% of all genes in the genome (N=16.434) had gonad-biased expression (N=4,242), and 9.6% of genes (N=1,573) had biased expression in the GT-soma (shown in Fig. S1). The N values of sex-biased genes for those genes with orthologs (N=7,751) are shown in Fig. S2 (N=2,341 (30.2%) and 836 (10.7%) for gonads and GT-soma respectively). We then assessed the sex-biased expression status of X-linked and autosomal genes with respect to dN/dS.

The proportion of genes on the X-chromosome and on each of the nine autosomes that had sex-biased or unbiased expression is shown in Fig. 2A, which includes all genes for which we had calculated dN/dS values (N=7,751) (see Fig. S3 for all annotated *T. castaneum* genes, which yielded similar patterns according to sex-biased expression status). We found that a disproportionately large fraction of genes on the X chromosome were ovary-biased: 53.9% of the X-linked genes under study were ovary-biased (N=233 of the 432 X-linked genes for which we assessed dN/dS) (Fig. 2A), while only 16.3% of autosomal genes showed ovary-biased expression (N=1,192 of 7,319 genes pooled across autosomes, Chi^2^ with Yate’s correction P<0.0001). In contrast, relatively few testis-biased, GT-male biased or GT-female biased genes were located on the X chromosome (each of these gene expression categories constituted ≤5.5% of the X-linked genes under study in Fig. 2AB). These chromosomal distributions of the different sex-bias expression categories for this set of 7,751 genes with high confidence orthologs between *T. castaneum* and *T. freemani* (Fig. 2AB) largely parallels that observed for all *T. castaneum* genes in the genome (Fig. S3AB) and agrees with the aforementioned prior report for all *T. castaneum* genes [38]. That study compared whole males versus whole females, and showed that the X-chromosome contained a high abundance of female-biased genes and very few male-biased genes. Our results extends these results to explicitly show that ovary-biased genes (Fig. 2A), rather than genes with female-biased expression in somatic tissues (Fig. 2B), are highly concentrated on the X-chromosome, and that X-linked testis-biased genes, GT-male-biased, and GT-female-biased genes are each relatively rarely observed on the X-chromosome.

**Fig. 2.**
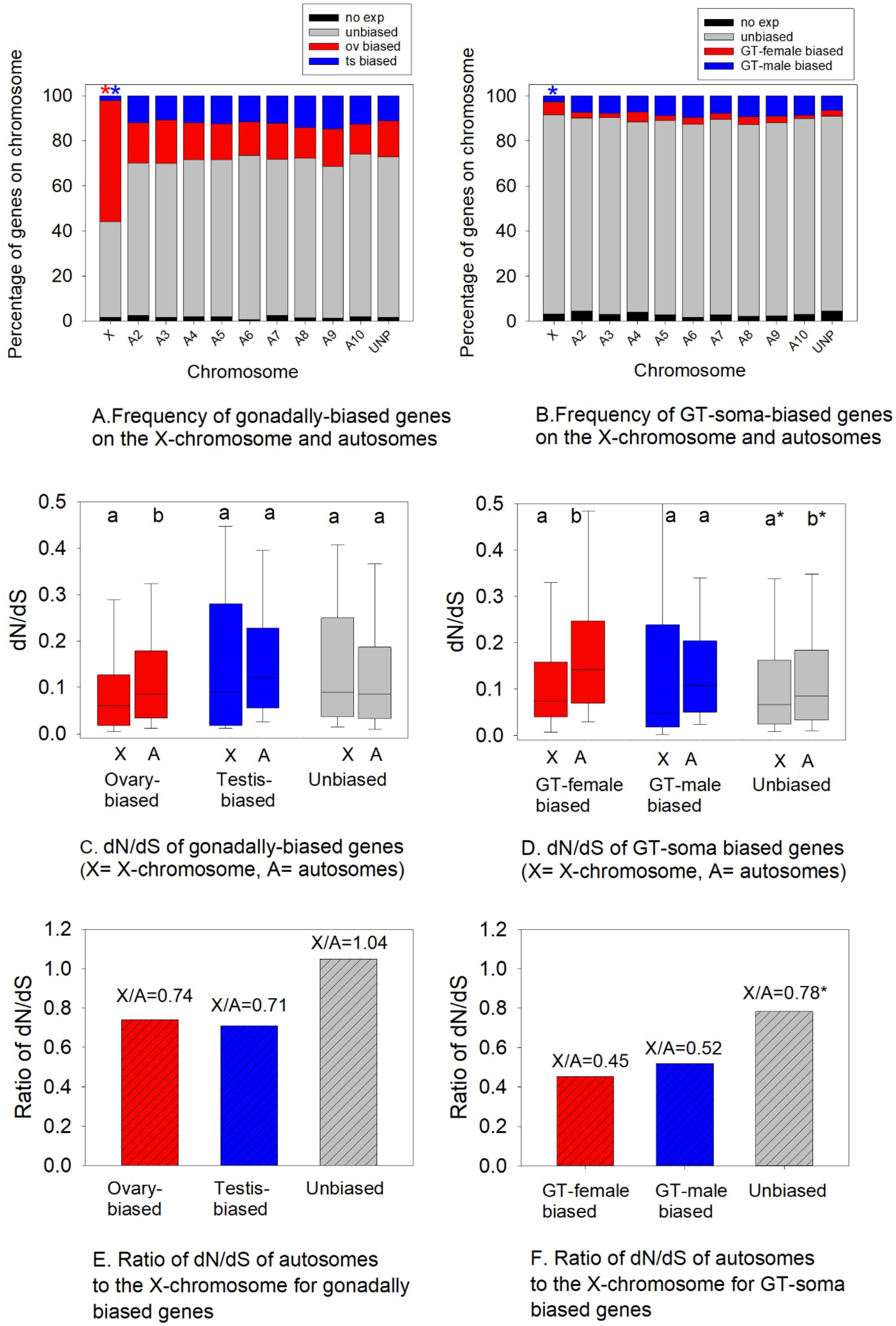
Assessment of the faster-X effect with respect to sex-biased genes in *T. castaneum.* A) The frequency of gonadally sex-biased genes on the X chromosome and nine autosomes for the 7,751 genes under study; B) the frequency for GT-soma sex-biased genes; C) the dN/dS of ovary-biased, testis-biased and unbiased genes on the X-chromosome and autosomes; D) the dN/dS of GT-male biased, GT-female biased, and GT-unbiased genes on the X-chromosome and autosomes; E) the ratio of the median dN/dS of the X chromosome to the autosomes (X/A_dN/dS_) for all three categories of sex-biased expression for the gonads; and F) for the GT-soma. In A, the red and blue asterisks indicate more ovary-biased and fewer testis-biased (or GT-male biased in B) genes were located on the X-chromosomes than on pooled autosomes (Chi^2^-P with Yate’s correction P<0.05 for each contrast). Different lowercase letters on top of each pair of bars in C and D indicate MWU-test P<0.05. In C-F, unmapped genes were included with autosomal genes and their inclusion in or exclusion from the analysis yielded similar results. Unbiased genes included those with no detectable expression. *Note that differences in X-linked and autosomal unbiased genes in panel D and F are explained by ovary-biased genes (Table S2) as outlined in the main text. After removal of ovary-biased genes X/A_dN/dS_=1.04 for GT-unbiased genes.

### The absence of a faster-X effect is largely caused by slow-evolving X-linked ovary-biased genes

Having identified that ovary-biased genes were highly overrepresented on the X chromosome relative to autosomes (Fig. 2A), we asked if this might contribute to the observed slower-X effect. We compared dN/dS values for these ovary-biased genes on X-chromosomes to those values for autosomal ovary-biased genes (Fig. 2CE; N values in Table S2, Fig. S2). We found that the dN/dS values of X-linked ovary-biased genes were statistically significantly lower than dN/dS values for autosomal ovary-biased genes (MWU-test P<0.001, Fig. 2C). Thus, the faster-X effect is absent in ovary-biased genes. Further, the ratio of the median dN/dS values when calculated using only the subset of X-linked ovary-biased genes versus those on autosomes, X/A_dN/dS (ovary-biased)_, was 0.74 (Fig.2E), also suggesting higher selective constraint on ovary-biased genes on the X-chromosome than autosomes. Moreover, ovary-biased genes on the X-chromosome had lower dN/dS than unbiased genes on the X-chromosome and on autosomes (MWU-tests P<0.001) and markedly lower dN/dS values than testis-biased genes on the autosomes (two-fold lower, 0.060 versus 0.120 median dN/dS values, MWU-test P<0.001; note there were too few X-linked testis-biased genes for reliable testing). Together, given the high frequency of genes located on the X-chromosome that were ovary-biased (Fig. 2A), these findings indicate that constrained evolution of ovary-biased genes contributes to the global absence of a faster-X effect in this organism (Fig. 1).

For the genes with GT-soma-biased expression, there were only 24 genes with GT-female biased expression on the X-chromosome (as compared to 233 with ovary-biased expression on the X-chromosome). Nonetheless, as we had observed for ovary-biased genes, this small number of GT-female biased genes also had statistically significantly lower dN/dS values than the GT-female biased genes on autosomes (MWU-test P=0.031, Fig. 2D), and the X/A_dN/dS (GT-female)_ value when calculated for this subset of genes was also low, at 0.45 (Fig. 2F). Thus, it appears that there has also been high purifying selection on X-linked GT-female biased genes in this taxon. Upon close examination however, and as shown in Table S2, 17 of the 24 (70.8%) X-linked GT-female biased genes also had ovary-biased expression, suggesting that the observed effect could be due to purifying selection on ovarian expression rather than somatic expression. Nonetheless, the seven genes with GT-female biased but not ovary-biased expression yielded a X/A_dN/dS (GT-female)_ ratio of 0.32, indicating that X-linked GT-female-biased genes are under higher constraint than those on autosomes, regardless of their ovary-biased expression status. Thus, we find no evidence of a faster-X effect for any female-biased genes, regardless of gonadal or somatic expression, and in fact these genes likely contribute to the slow evolution of the X-chromosome.

We next assessed whether the faster-X effect was observable for male-biased genes (testis- or GT-male-biased), which would be expected to exhibit a pronounced faster-X effect under a hypothesis of rapid fixation of beneficial recessive mutations in the heterogametic sex [7, 16]. We found that very few testis-biased genes or GT-male-biased genes were located on the X chromosome (N=9 and N=12 for testis-biased and GT-male-biased X-linked genes with high confidence interspecies orthologs), and that neither group of male-biased genes showed even mild evidence of a faster-X effect. The median dN/dS value was lower for these genes on the X chromosome than on autosomes for both categories of genes (Fig. 2CD). The X/A_dN/dS (testis-biased)_ ratio was 0.71 for testis-biased genes, and the X/A_dN/dS (GT-male biased)_ ratio was 0.52 for GT-male biased genes (Fig. 2EF), markedly below 1 in both cases. No overlap was observed between the testis-biased and GT-male biased gene sets (Table S2), and thus the low dN/dS effects were independently observed in each group. For stringency, we examined and noted that three of the GT-male-biased genes were also ovary-biased, but exclusion of those genes from the analysis still yielded an X/A_dN/dS (GT-male biased)_ ratio of 0.59, and thus the low dN/dS effect is directly linked to the GT-male-biased expression. In sum, while the small number of X-linked testis-biased and GT-male-biased genes precludes rigorous statistical testing of those genes, the patterns observed for these genes are inconsistent with a faster-X effect in male-biased genes, whether gonad- or soma-biased.

We next asked whether there was evidence for the faster-X effect in the gonadally unbiased genes. Given that such genes were common on all chromosomes (Fig. 2A, Table S2), which provides the potential for high statistical power, and that they by definition they exclude the highly constrained X-linked ovary-biased genes and the testis-biased genes described above (Fig. 2C), we predicted that if there were even a mild tendency for a faster-X effect in this taxon, it would be readily apparent in this group of genes. However, we found no significant difference in dN/dS values between X-linked and autosomal gonadally unbiased genes (MWU-test P>0.05 Fig. 2C). Rather, we observed an X/A_dN/dS (gonadally unbiased)_ ratio of 1.04, indicating highly similar dN/dS between these two groups (Fig. 2E). In this regard, we conclude that the faster-X is fully absent in gonadally unbiased genes.

Finally, we assessed the GT-unbiased genes, and found evidence for greater constraint on the sequence evolution of these genes on the X chromosome as compared to autosomes (X/A_dN/dS (GT-unbiased)_ =0.78, MWU-test P<0.05, Fig. 2DF). As expected, however, given that a majority of X-linked genes under study were ovary-biased (Fig. 2A, Table S2), and that most genes expressed in the GT-soma are not sex-biased (Fig. 2B), many of the X-linked GT-unbiased genes (N=396) were also ovary-biased (N=213). Excluding these genes, so that we could consider only those 183 GT-unbiased genes that were not ovary-biased, we found no differences in dN/dS values for these genes between the X-chromosome and autosomes (MWU-test P>0.05). In fact, the X/A_dN/dS (GT-unbiased)_ ratio for these GT- and ovary-unbiased genes was 1.04, identical to that observed for gonadally unbiased genes (Fig. 2EF). Thus, the GT-somatically unbiased genes, whether they were co-biased in the ovaries or not, exhibited no signals of a faster-X effect.

Taken together, the collective results in Fig. 2 show that the slower-X effect observed here in *Tribolium* is largely explained by highly constrained evolution of the abundant X-linked ovary-biased genes, with some minor contributions from the relatively smaller number of testis-biased, GT-male biased, and GT-female-biased genes (Fig. 2C-F). Crucially, the faster-X effect was not even observed in either gonadally-unbiased or GT-soma-unbiased genes, which each yielded an effective X/A_dN/dS_ ratio of 1.04. This latter finding cannot be explained by constrained evolution of X-linked sex-biased genes, suggesting that other factors likely also contribute towards the absence of the faster-X in this taxon (see the below section “*Absence of dosage compensation in the T. castaneum testis*”).

### Why do X-linked ovary-biased genes evolve slowly?

We wished to further consider why the X-linked ovary-biased genes evolved extremely slowly (Fig. 2CE). The exceptionally low dN/dS values observed for ovary-biased genes on the X chromosome (Fig. 2CE) as compared to autosomes suggests that they could be essential genes subjected to high purifying selection, and their ovary-biased expression suggests that they may be involved in female reproduction and thus fitness. To examine this, we determined the predicted GO functions (see Methods: GO functions determined in DAVID [45]) of the ovary-biased genes located on the X-chromosome (Fig. 2A). Indeed, in agreement with this hypothesis, we found that ovary-biased genes on the X chromosome were enriched for genes involved in ovarian follicle development and *wnt* signalling (Table 1), which is crucial for ovarian development and function in multiple animals (see [46–55] for examples). X-linked ovary-biased genes also included those with predicted roles in female meiosis and oocyte function (Table 1). These essential ovarian roles were not among the top functional categories observed for ovary-biased genes on autosomes (Table 1). Given these results, we suggest that high purifying selection on ovary-biased genes on the X chromosome is likely at least partly due to the important female reproductive roles of some of these genes. Moreover, their high concentration on the X-chromosome may suggest a history of preferential translocation of essential female reproductive genes to the X-chromosome.

**Table 1.** Gene ontology (GO) clustering of ovary-biased genes located on the X chromosomes and on autosomes. The top clusters with the greatest enrichment scores are shown per category. *P*-values are from a modified Fisher’s test, wherein lower values indicate greater enrichment. Data is from DAVID software [45] using those genes with *D. melanogaster* orthologs.

| Ovary-Biased Genes on X Chromosome |  | Ovary-Biased Genes on Autosomes<sup>a</sup> |  |
| --- | --- | --- | --- |
| **<u>Cluster 1: Enrichment Score 3.09</u>** | **P-value** | **<u>Cluster 1: Enrichment Score 3.56</u>** | **P-value** |
| Wnt signaling pathway | 4.20E-06 | Metal-binding | 6.00E-05 |
| Segmentation polarity protein | 8.20E-05 | Zinc ion binding | 5.50E-04 |
| Regulation of Wnt signaling pathway | 1.60E-04 | Zinc-finger | 6.60E-04 |
| Segment polarity determination | 1.30E-03 |  |  |
| Ovarian follicle cell development | 6.70E-03 | **<u>Cluster 2: Enrichment Score 2.81</u>** |  |
| Somatic stem cell population maintenance | 2.50E-02 | Pleckstrin homology-like domain, signalling | 1.80E-04 |
| Heart development | 3.90E-02 | Pleckstrin homology domain, signalling | 4.90E-04 |
| **<u>Cluster 2: Enrichment Score 2.92</u>** |  | **<u>Cluster 3: Enrichment Score: 2.78</u>** |  |
| ATP-binding | 2.00E-04 | SH2 domain, oncoproteins, signalling | 1.40E-04 |
| Nucleotide-binding | 3.70E-04 | SH3 domain, intracellular or membrane-associated proteins | 2.10E-04 |
| Nucleotide phosphate-binding region:ATP | 1.60E-03 |  |  |
| Protein kinase, ATP binding site | 7.70E-03 |  |  |

<sup>a</sup> Data was pooled for all nine autosomes and also includes genes yet unmapped in the genome.

We next considered whether expression breadth could explain the slow evolution of X-linked ovary-biased genes. It has been proposed that greater expression breadth across tissues, which reflects pleiotropic functionality, constrains dN/dS. For example, the rapid evolution of male-biased than female-biased genes observed in various organisms, as was also found here for testis versus ovaries (MWU-test P<0.001 of all testis-versus all ovary-biased genes, Fig. 2C) may result from low pleiotropy [17, 40, 56, 57]. Indeed, we found herein that expression breadth across the four studied tissue-types was lower for testis-biased than for ovary-biased genes. Specifically, only 25.5% of testis-biased genes (pooled for X-linked and autosomal) were expressed in all four tissue types (at >1FKPM) while 72.8% of ovary-biased genes were transcribed in all four tissues. In this regard, ovary-biased genes as a group exhibit higher pleiotropy, suggesting potential roles across various tissues that may contribute to their slower evolution relative to testis-biased genes (Fig. 2C). In turn, the accumulation of ovary-biased genes on the X-chromosome would act to constrain evolution of this chromosome.

Nonetheless, it is worth noting that broad expression breadth (expressed in in all four tissues) was observed for the majority of ovary-biased genes independently of chromosomal location (78.9% of X-linked ovary-biased genes and 71.6% of autosomal ovary-biased genes were expressed in all tissues). Thus, the specific finding of a lower dN/dS values of X-linked ovary-biased genes (compared to their counterparts on autosomes, Fig. 2C) cannot be fully explained by high pleiotropy. We therefore propose that the slower evolution of ovary-biased genes on the X-chromosome than those on autosomes is likely at least partly due to their preferential involvement in essential ovary functions (Table 1).

### Absence of dosage compensation in the *T. castaneum* testis

In X/Y sex determination systems, it has been posited that mechanisms should exist to ensure that the expression levels of genes on the X-chromosome (X) and autosomes (A) would be approximately equivalent in both males (with hemizygous X) and females (homozygous X), such that the ratio of expression of X/A in each sex should equal one [38, 58]. In turn, it may be expected that X_male_/X_female_ = A_male_/A_female_ = 1 [38]. Measurements of gene expression levels in a number of animals have shown that mechanisms for acquiring elevated expression on the single male X-chromosome, or dosage compensation, are highly variable and that full dosage compensation is sometimes, but not always, achieved [38, 58–60]. In one prior study of gene expression using microarrays of whole males versus whole females in *T. castaneum* [38], it was reported that males exhibited full X-chromosome dosage compensation, with X_male_/A_male_ = 1.0 and that females exhibited overexpression of the X chromosome, with X_female_/A_female_ = 1.5, thereby yielding X_male_/X_female_=0.79 and A_male_/A_female_ =1. Those results were interpreted as evidence that the genes on the X-chromosome exhibited complete dosage compensation in males (meaning that expression of the hemizygous X linked genes was equalized to expression of autosomal genes in males), and were overexpressed in females as an imperfect response to dosage compensation [38]. However, a recent study that examined published RNA-seq data for somatic glandular tissues in *T. castaneum* did not find evidence for hypertranscription of the X-chromosome in females [60]. Given that dosage compensation can vary among tissues in insects, particularly its absence in the testis of *Drosophila* [25–27], and that complete dosage compensation has been theorized to promote the faster-X effect by fixation of recessive beneficial mutations in hemizygous males [1, 2, 22], we next aimed to assess dosage compensation separately for genes expressed in the gonads and those expressed in the GT-somatic tissues for *T. castaneum*. As the sex organs play central roles in reproduction, recessive beneficial mutations in genes are apt to have their greatest fitness consequences (and thus be fixed) in the hemizygous male gonad (rather than male soma), and thus we predicted that if a lack of dosage compensation were found in the testis, this might significantly contribute towards the absent faster-X in this organism.

In Fig. 3, we show the median expression level (FPKM) for genes on the X-chromosome and each of the nine *T. castaneum* autosomes for the gonads (A) and for the GT-soma (B) using all genes that had high-confidence *T. freemani* orthologs (N=7,751; results for all *T. castaneum* genes are in Fig. S4, showing similar patterns). We report that expression levels in ovaries (Ov) were largely similar across the nine autosomes (median 14.7 FPKM across nine autosomal medians) and were relatively elevated on the X-chromosome (18.8 FPKM, MWU-test P=0.023 of the X-chromosome versus autosomes, Fig. 3A; note that X/A is measured using multiple decimal places), yielding X_OV_/A_Ov_ of 1.26 and is consistent with overexpression of X-linked genes in the ovary. For the testis (Ts), however, while expression was also largely similar across all nine autosomes (median 7.9 FPKM across nine autosomal medians), a strikingly lower expression level was observed for the X-chromosome (3.2 FPKM, Fig. 3A), giving an X_Ts_/A_Ts_ value of 0.41. Thus, there is 2.5-fold lower expression of X-linked testis genes than of autosomal testis genes (MWU-test P<0.001, Fig 3A; see also Fig. S4A where the value was also <0.5), inconsistent with hypertranscription of the single X chromosome in males, at least for the testis-expressed genes. This complete absence of dosage compensation in the *T. castaneum* testis is even beyond that reported for the testis of *Drosophila*, which had an 0.65 value for this parameter [25]. Further, the low value potentially not only suggests an absence of hyperexpression on the X-chromosome in the hemizygous state (to balance autosomes), but could also be consistent with an active mechanism of suppression of X-linked expression [25, 27, 61] in this beetle.

**Fig. 3.**
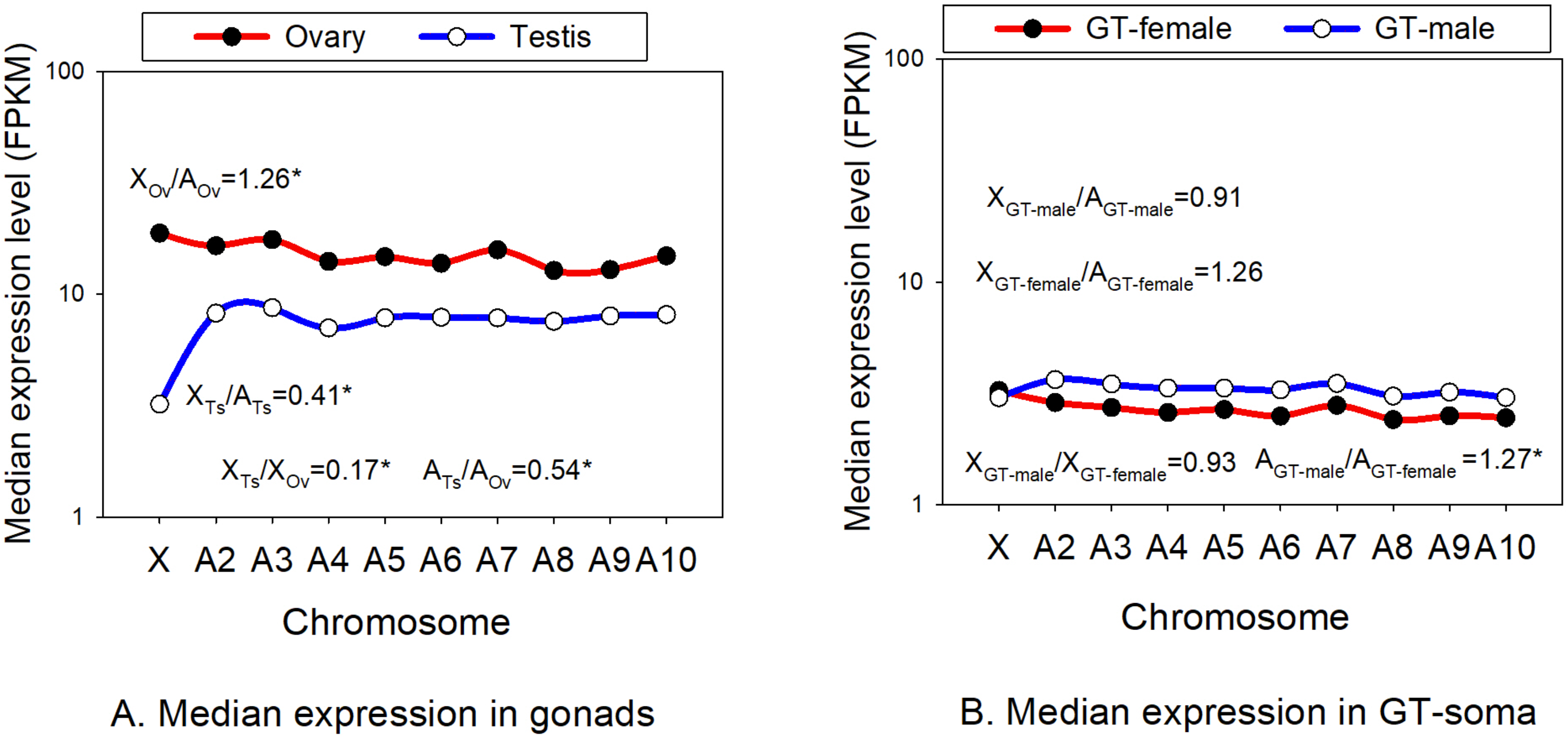
Median expression in the male and female tissues on each of the ten chromosomes in *T. castaneum* for all genes under study. A) Gonads; B) GT-soma. For panel A, the ratio of median expression on the X chromosome (X) and autosomes (A) for testis-biased genes and for ovary-biased genes are shown (X_Ts_/A_Ts_ and X_Ov_/A_Ov_). Also shown are X_Ts_/X_Ov_ and A_Ts_/A_Ov_. Panel B contains the equivalent results for the GT-soma. *Indicates a statistically significant difference between the two groups contained in each ratio using MWU-tests.

Moreover, we found that testis expression was lower than ovary expression across all nine autosomes, such that A_Ts_/A_Ov_ was equal to 0.53 (MWU-test P<0.001 of autosomal testis to ovary expression), differing from the equal male/female expression typically expected on autosomes [25, 38]. This effect was even more pronounced for the X-chromosome, where X_Ts_/X_Ov_ had a value of 0.17 (Fig. 3A, MWU-test P<0.001 for X-chromosome testis expression versus ovary expression), indicating that even after taking into account the lower expression level observed on all autosomes for testis genes versus ovary genes (median 1.9 fold), testis genes exhibited a marked drop (5.9-fold) in expression on the X-chromosome. In this regard, both X_Ts_/A_Ts_ and X_Ts_/X_ov_ (Fig. 3A) suggest a complete absence of dosage compensation in this beetle.

Considering the GT-soma, we observed nearly perfect dosage compensation on the X-chromosome for GT-males, both with respect to GT-female expression levels, such that X_GT-female_/X_GT-male_ =0.93 (median of 3.02 and 3.25 FPKM respectively MWU-test P=0.74), and with respect to autosomal GT-male expression levels, with X_GT-male_/A_GT-male_=0.91 (MWU-test P=0.11). Thus, unlike genes expressed in the testis, genes expressed in the non-gonadal tissues of males (GT-males) exhibited high dosage compensation (Fig. 3B, Fig. S4B). The median GT-male expression across all nine autosomes was consistently higher than the median expression in GT-females, yielding A_GT-male_/A_GT-female_ of 1.27 (MWU-test P<0.001), a trend opposite to the higher expression level observed for ovary genes relative to testis genes (Fig. 3AB). Nonetheless, GT-female genes on the X-chromosome were expressed at higher levels than such genes on autosomes, yielding X_GT-female_/A_GT-female_=1.26 (MWU-test P=0.064), and thus contributing to the observed highly similar expression levels between GT-females and GT-males on the X-chromosome. In sum, the GT-males show evidence of nearly complete dosage compensation, differing markedly from its complete absence in the testis. Additional study of more individual somatic tissues (e.g., brain, hindgut), similar to that in other recent studies [25, 60], will be needed to assess whether the variation in GT-female expression among autosomes is observed in various somatic tissue types in *T. castaneum*.

Taken together, the results presented here in Fig. 3 and Fig. S4 show a complete lack of dosage compensation in the testis. Given that the faster-X effect may be anticipated to be strongest in taxon groups with complete dosage compensation, due to elevated phenotypic protein product and effects of beneficial recessive mutations in males [1, 14, 22], the under-transcription in the *Tribolium* testis could contribute to the absence of the faster-X we report here (Fig. 1, Fig. 2). Further, this effect may transcend all sex-biased expression categories. For instance, all X-linked testis-biased genes, 98.7% ovary-biased genes, and 92.3% of gonadally unbiased genes were expressed in the testis, and thus low dosage compensation may affect all groups of genes due to under-expression (relative to autosomal genes) on the single X-chromosome in males. Finally, it should be noted that the absence of X-chromosome dosage compensation found in the testis, combined with a relatively modest elevation in ovary expression (Fig. 3A), are consistent with the concentration of ovary-biased genes on the X chromosome in this organism (Fig. 2AB, Fig.S3AB).

## Discussion

### Absence of a faster-X effect and sex-biased genes

Our results show that the absence of a faster-X effect, and tendency for a slower-X, in *Tribolium* (Fig. 1), is explained in part by strong purifying selection on the highly abundant X-linked ovary-biased genes (Fig. 2ACE). Accordingly, we hypothesize that many X-linked ovary-biased genes play essential roles in this organism, as indicated by their low dN/dS values (Fig. 2CE), GO functional analysis (Table 1) and high cross-tissue pleiotropy (72.8% expressed in all four tissues). We further hypothesize that ovary-biased genes may have been preferentially translocated to the X-chromosome over the history of this beetle taxon. This type of localization, or translocation, phenomenon is also supported by findings from other systems (e.g. mice, humans, flies) showing that genes involved in sex and sexual dimorphism, and particularly those showing female-biased expression, have been preferentially localized to the X-chromosome [62–65]. The preferential transfer of female genes (as compared to male-biased genes) to the X-chromosome may be facilitated by the unique selection pressures experienced by these chromosomes, especially the fact that two-thirds of X-chromosomes in population are carried by females and only one third in males, which may make the concentration of female functional genes on the X-chromosome an innate benefit to females [18, 61, 65].

For the testis-biased and GT-male biased genes, of which there were very few on the X-chromosome (Fig. 2AB, Fig. S3AB), the low dN/dS observed on the X-chromosome (X/A_dN/dS_=0.71 and 0.52 respectively) is also discordant with a faster-X effect in those genes. The constraint on X-linked male-biased genes could be readily explained by the immediate exposure of any mildly deleterious recessive mutations to purifying selection in the hemizygous state in testis and GT-males (and not on autosomes). Thus, it is possible that a much different mechanism could cause the slow evolution of those genes, than that operating on ovary-biased genes. In other words, it may be surmised that if most protein sequence evolution is due to fixation of weakly deleterious alleles in beetles, then the X-chromosome would be expected to evolve slowly due to rapid purging of these mutations as a result of hemizygous exposure, as compared to autosomes. This phenomenon was theorized to occur for some organisms under the original faster-X hypothesis [1] and has been proposed to contribute to the relatively mild faster-X effect observed in *Drosophila* (by countering the accelerated evolutionary rates on the X due to rapid fixation of recessive beneficial mutations) [22]. In Satyrine butterflies with W/Z systems, for example, it was suggested that slow evolution of Z-linked female-biased genes occurred due to high purifying selection on the Z-chromosome in the hemizygous state [13]. Thus, the same phenomenon may occur for male-biased genes in *Tribolium*.

Importantly, the unbiased genes in the gonads, and in the GT-somatic tissues, showed no tendency for a faster-X effect, with X/A_dN/dS (unbiased)_ =1.04 for each tissue type (after exclusion of ovary-biased genes from the latter dataset, see Results, Fig. 2C-F). Thus, the lack of faster-X effect in those genes cannot be explained by higher selective constraint on the X-chromosome than autosomes. Albeit, we do not exclude some purging of recessive mutations in males could occur in these genes, similar as proposed possible for male-biased genes above, however this would be expected to slow the X [1], which was not observed for unbiased genes. The result indicates that another mechanism likely contributes to the absence of a faster-X effect in these beetles, which our data strongly suggest involves the lack of dosage compensation in the male gonads.

### Lack of dosage compensation

Our finding of a complete absence of dosage compensation combined with the absence of the faster-X effect is highly suggestive that fixation of beneficial sequence mutations on the X-chromosome may have been uncommon or absent in this taxon due to under-expression in males [1, 22]. Additional findings of an absent faster-X effect in a broader range of organisms, and that include assessments of the degree of dosage compensation, will help further resolve the role of this putative mechanism. Our results suggest the effect may extend beyond the passive absence of dosage compensation (or hyperexpression of the X-chromosome), and potentially reflects an active mechanism of X-chromosome silencing. Recent reports from *Drosophila* have also shown that dosage compensation is absent for the testis [25], and that this insect exhibits active suppression of X-linked expression in males [25, 61]. Consistently, it was found that the transfer of X-linked testis-expressed genes to the autosomes resulted in marked upregulation in *D. melanogaster* [61], suggesting an active mechanism of suppression of expression on the X-chromosome in testis. While the mechanism for X-linked active suppression of expression is unknown, it could reflect male meiotic sex chromosome inactivation (MSCI). Empirical support for MSCI has been observed for *D. melanogaster* [27, 66], and a strong effect has been found in range of other animal systems including mammals [67] and *Caenorhabditis elegans* [68]. Further study will be needed to ascertain whether the absence of dosage compensation in the testes for *T. castaneum* involves lack of upregulation on the X-chromosome and/or also includes an active process involving X-chromosome suppression or silencing.

We emphasize that the separation of gonads and GT-somatic tissues herein for expression analyses was essential for revealing the absence of dosage compensation in *T. castaneum*. A prior report of dosage compensation in *T. castaneum* using microarray expression data from whole males and females [38] showed much different results for male dosage compensation than those observed in the present study. In that assessment, males were reported to exhibit complete dosage compensation on the single X-chromosome as compared to autosomes, while females were reported as exhibiting overexpression of X-chromosomes (relative both to X-linked transcription in males and to female expression on autosomes) [38]. The latter was interpreted as an imperfect (female) response to male dosage compensation [38]. In contrast, our tissue-specific expression data allow us to explicitly show the absence of dosage compensation in the male testis, and nearly perfect dosage compensation for X-linked GT-male expression with respect to the autosomes and with respect to X-linked expression in GT-females. Thus, our separation of gonadal from somatic expression data was essential for the detection of key differences in dosage compensation in this taxon, and was particularly crucial to discerning the absence of dosage compensation in the testis (Fig. 3A, Fig. S4A), that had been obscured previously in the examination of whole males and females.

Finally, while we propose that the absence of the faster-X effect herein is best explained by constrained evolution of the abundant X-linked ovary-biased genes and lack of dosage compensation in the testis, we do not exclude a role of standing genetic variation. For instance, large populations tend to contain more polymorphic loci, which can accelerate autosome evolution if adaptation occurs via standing genetic variation rather than *de novo* mutations [2, 69]. This phenomenon could possibly occur in beetles, and thus we do not exclude this factor in partly contributing towards the absence of a faster-X effect in this taxon.

## Conclusions and Future Directions

We have shown the complete absence of the faster-X effect in a *Tribolium* system, which our data strongly suggest is largely explained by constrained evolution of X-linked ovary-biased genes and the extreme absence of dosage compensation in the testis of this taxon. Future studies should aim to study additional genomes of more *Tribolium* species, which would allow tests of positive selection in protein sequences on the X-chromosome and autosomes [43]. In addition, studies using population-level data from *T. castaneum* will allow tests of polymorphism versus divergence, which comprises an alternate method to test adaptive evolution expected under the faster-X effect [13, 14]. A further understanding of dosage compensation in this taxon may be achieved by attainment of transcriptional data from a wide range of individual somatic tissue types in *T. castaneum*, similar to analyses recently conducted in *Drosophila* [25]. Such multi-tissue expression data will also allow further assessments of cross-tissue pleiotropy of sex-biased genes [40, 57, 70] and may help further disentangle its role in constraining evolution of ovary-biased genes (Fig. 2).

Moreover, experimental research of MSCI in *T. castaneum,* as has been conducted in other organisms [66, 67, 71], will help reveal whether the lack of dosage compensation observed in the testis is due to transcriptional silencing in the male meiotic cells. In addition, studies using X-linked genes inserted into the autosomes, and vice-versa [25, 61, 68] may help discern the dynamics of dosage compensation in *T. castaneum*. Finally, studies of the faster-X effect, including analyses of sex-biased genes and dosage compensation, should be extended to include more understudied organisms, to help reveal the breadth of this phenomenon in metazoans and to better understand its underlying mechanisms.

## Methods

### Biological Samples and RNA-seq

*T. castaneum* and *T. freemani* specimens were provided by the Brown lab at Kansas State University (strain IDs; https://www.k-state.edu/biology/people/tenure/brown/). Samples were grown under standard laboratory conditions until adulthood as previously described [29]. Additionally, to ensure that all adult animals for both species remained unmated until the time of tissue dissection, all animals were separated as late stage larvae into individual vials containing flour and allowed to pupate into adults. Tissue dissections were then performed on unmated adults within a week after they emerged from the molt. For *T. castaneum* a total of 150 animals per sex per biological replicate were sampled, whereas for *T. freemani*, a total of 50 males and females were collected per sample. For each sample of males and of females, the gonadal and nongonadal tissues were separated and placed into two separate vials containing TRIzol reagent (Ambion Life Technologies, catalog number 15596-018) on dry ice. Technical details on tissue collection, PCR, and RNA-seq are provided in Additional File 1, Text File S1.

For males, the isolated reproductive tissues included the testes, accessory glands (mesadenia, ectadenia), and directly attached tissues (vesicular seminalis, vas deferens and ejaculatory duct) whilst for females, gonad samples included the ovaries and their linked tissues (spermathecal gland, common oviduct, spermathecae, and vagina). For simplicity, we refer to the male and female reproductive organs and tissues collectively as “testis” and “ovary” or the sex-neutral “gonads” herein, with the understanding that they include the abovementioned reproductive tissues directly linked to the respective gonads. All remaining non-gonadal tissues of the adult body are referred to as the gonadectomized (GT-) soma, or GT-males and GT-females. For the sister species *T. freemani*, four RNA-seq samples, one per tissue-type, testes, ovaries, GT-males and GT-females, were obtained and used for refining the CDS list for this species (see Methods) that was employed to assess protein divergence (dN/dS).

### CDS per Species and Defining Orthologs

The annotated CDS of our main target species *T. castaneum* (v.5.2) were downloaded from Ensembl Metazoa (http://metazoa.ensembl.org) and are also available at BeetleBase [34, 36]). The full CDS per gene (longest CDS per gene) was used for the study of sex-biased gene expression.

For the genome of *T. freemani*, which we used as a reference to determine dN/dS, CDS have not yet been annotated and thus were extracted from available scaffolds. The scaffold assembly was downloaded from BeetleBase (version 4, http://www.Beetlebase.org, [34]). Details on extracting the CDS for *T. freemani* are provided in Additional File 1, Text File S1.

In the final CDS list for *T. castaneum* and for *T. freemani*, only those CDS having a start codon, not having unknown or ambiguous nucleotides or internal stop codons, and ≥33 amino acids were retained for study. The total number of CDS after filtering was 16,434 for *T. castaneum*, marginally more than the 16,404 gene models first defined for this species [36], and was 12,628 for the sister species *T. freemani*. The average GC content of the *T. castaneum* protein-coding genes was 46.1% (±5X10^-4^), which is above the 33% reported for the global genome encompassing all coding and noncoding DNA as has been noted previously for this taxon [35, 36].

### Identification of Sex-Biased Genes

The RNA-seq reads (76bp) per sample were trimmed of adapters and poor-quality bases using the program BBduk available from the Joint Genome Institute (https://jgi.doe.gov/data-and-tools/bbtools/) and run as a plug-in in Geneious v11.0.3 using default parameters.

Gene expression level per gene was determined by mapping each RNA-seq dataset per tissue to the full CDS list for each species using Geneious Read Mapper, a program based our comparisons and other analyses provides similar read match performance as other common read-mappers such as Bowtie [72] or BBmap (https://jgi.doe.gov/; data not shown). Read counts per CDS were converted to FPKM for each gene. Expression level was compared separately for the gonads and for the GT-soma. Expression was compared between the testes and ovaries, and between GT-males and GT-females by using Deseq2 to obtain P-values [44] and the average FPKM of the replicates per tissue type (Table S1). Any gene having at least a two-fold difference in average expression and a statistically significant P-value (P<0.05) as well as a FPKM of at least one in one tissue type was identified as sex-biased [17, 73]. All other genes with nonzero expression in gonadal and in nongonadal contrasts were defined as unbiased.

### Ortholog Identification and Sequence Divergence

For dN/dS analysis, orthologs between *T. castaneum* and *T. freemani* were identified using reciprocal BLASTX of the full CDS list between species in the program BLAST+ v2.7.1 (https://blast.ncbi.nlm.nih.gov). Only genes having the same best match in both forward and reverse contrasts and an e-value <10^-6^ were defined as orthologs. In the rare cases when two CDS had the same e-value, the one with the highest bit score was taken as the best match. For additional stringency of genes used for the study of dN/dS, only those genes that were reciprocal BLASTX best matches and where both dN and dS values of alignments (≥33 amino acids) had values <1.5, and thus were unsaturated in substitution (see below paragraph), were defined as orthologs between *T. castaneum* and *T. freemani* (and used for dN/dS analyses). Thus, the alignments and dN/dS measures herein are conservative.

Orthologous gene sequences in *T. freemani* and *T. castaneum* were aligned by codon using MUSCLE set to default parameters (except the gap penalty which was set at −1.9) in the program Mega-CC v7 [74]. Alignments were then filtered to remove gaps. It has been found that removal of highly divergent segments from alignments, despite some loss of sequence regions, improves measures of protein divergence; thus, highly divergent segments were removed using the program Gblocks v. 0.91b set at default parameters [75, 76]. Each gene alignment was then run in yn00 of PAML, which accounts for codon usage biases [43], to measure dN, dS, and dN/dS [43].

We note that the percentage of ovary-biased genes on the X-chromosome in our 7,751 *T. casteneum* genes with orthologs in *T. freemani* was 53.9%, while for all 16,434 *T. castaneum* genes it was 42.2%. This likely reflects the fact that we studied genes that had high confidence orthologs between species, which are apt to be more frequently identified for ovary-biased genes due to their slowed evolution (Fig. 2C), an effect that was pronounced on the X-chromosome (Fig. 2C). Ovary-biased genes on autosomes had largely similar percentages on autosomes in the studied genes for dN/dS analyses and all genes (Fig. 2A, Fig. S3A). In this regard, our dN/dS analyses inherently include those genes with conserved between-species orthologs.

### X-Chromosomes Versus Autosomes

Chromosomal locations of genes are available in the annotation for *T. castaneum* (http://metazoa.ensembl.org, also available at BeetleBase [34, 36]). We note that the Y-chromosome of *T. castaneum* is small (<5MB), highly degenerate, contains few if any protein-coding genes, and is not included in the genetic linkage map; accordingly it was not studied [29, 36, 38, 77].

### Gene Ontology

Gene ontology (GO) was assessed using DAVID software [45]. For this, we identified orthologs to *T. castaneum* in the reference insect model *D. melanogaster* (CDS v6.24 available from www.flybase.org [78]) using BLASTX (https://blast.ncbi.nlm.nih.gov) to identify the best match (lowest e-value with cut off of e<10^-6^). *D. melanogaster* gene identifiers, which are accepted as input into DAVID, were used to obtain GO functions for *T. castaneum* genes. Single direction BLASTX with *T. castaenum* CDS as the query to the *D. melanogaster* database was used for this assessment (unlike for the reciprocal BLASTX between *Tribolium* species), as we considered reciprocal BLASTX to be overly stringent between these divergent insects (which are from different orders) for functional analysis.

### Availability of Data

The CDS v. 5.2 for *T. castaneum* are available at Ensembl Metazoa (http://metazoa.ensembl.org). Scaffolds for *T. freemani* are available at BeetleBase [34, 36]. RNA-seq data and SRA Biosample identifiers for all 12 samples from *T. castaneum* and *T. freemani* described in Table S1 are available at the SRA database under Bioproject accession number PRJNA564136.

## Declarations

### Ethics approval and consent to participate

N/A

### Consent for publication

N/A

### Availability of data and material

Available at the SRA database under Bioproject accession number PRJNA564136. See also Table S1 for SRA Biosample identifiers for each sample studied herein.

### Authors’ contributions

CAW, AK, and CGE designed the study, CAW conducted data analyses, AK conducted lab procedures, all authors contributed to, read and approved the final manuscript.

## Acknowledgments

This work was supported by funds from Harvard University. The authors thank Prof. Sue Brown at KSU for generously providing samples of *T. castaneum* and *T. freemani* for this study. We also thank members of the Extavour lab, Dr. Katharina Hoff at the University of Greifswald for updating the Augustus database used for *T. freemani* at our request, and the Bauer core sequencing facility at Harvard for generating RNA-seq data.

## Additional Files

Additional File 1: contains the supplemental tables, figures and SI Text including detailed methods.

## ADDITIONAL FILE 1

**Fig. S1.**
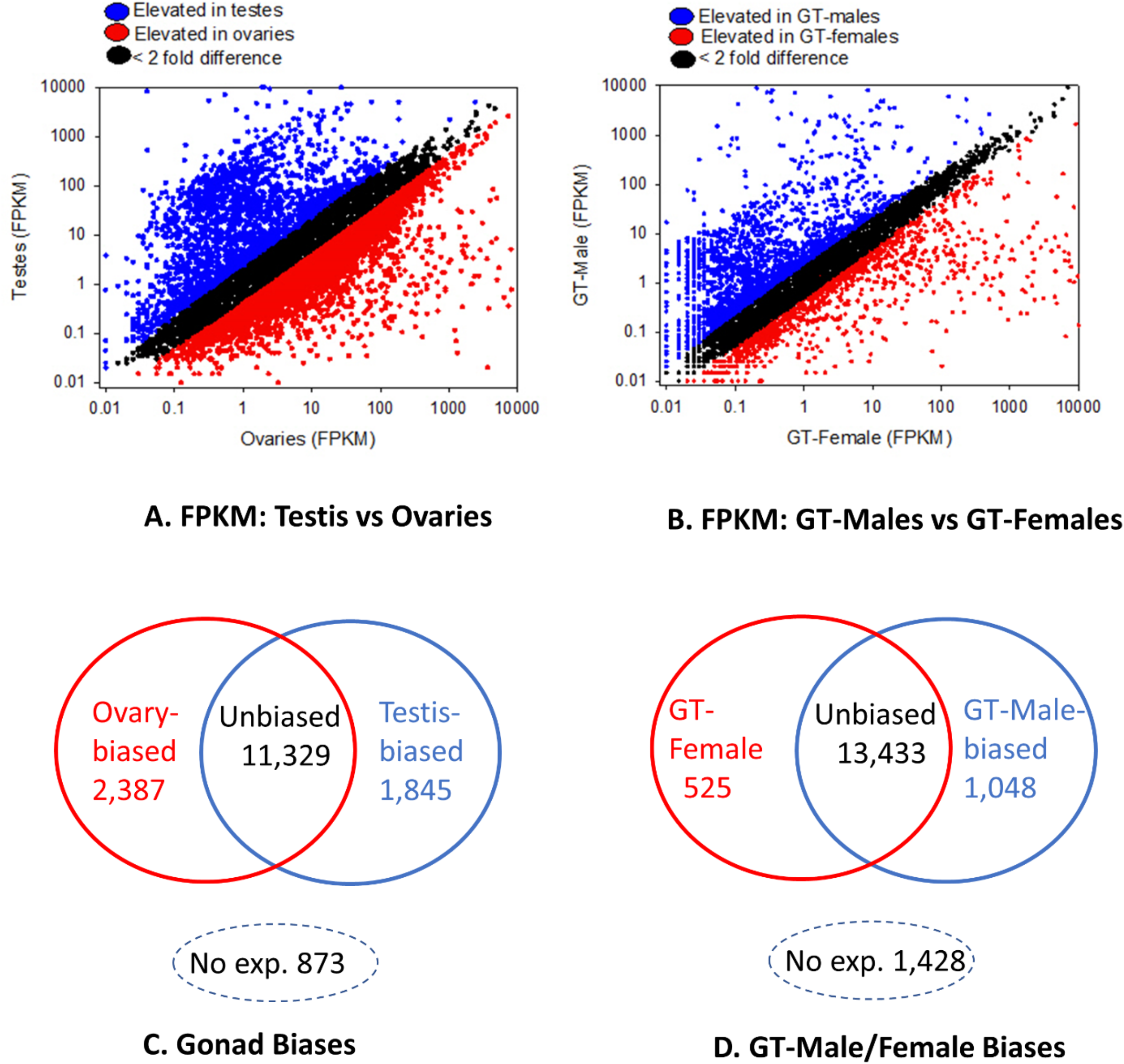
Sex-biased expression in 16,434 genes of *T. castaneum*. A) Expression level (FPKM) in the testes versus ovaries; B) expression level (FPKM) in GT-males versus GT-females; C) Venn diagram of sex-biased expression the gonads and; D) Venn diagram of sex-biased gene expression in the GT-soma. In A and B, all genes are shown and those with two-fold or greater difference in expression in the male and female tissues are in blue and red respectively (those statistically significant in C and D). Genes with no expression in both tissues were excluded in A and B and are shown in C and D (No exp.).

**Fig. S2.**
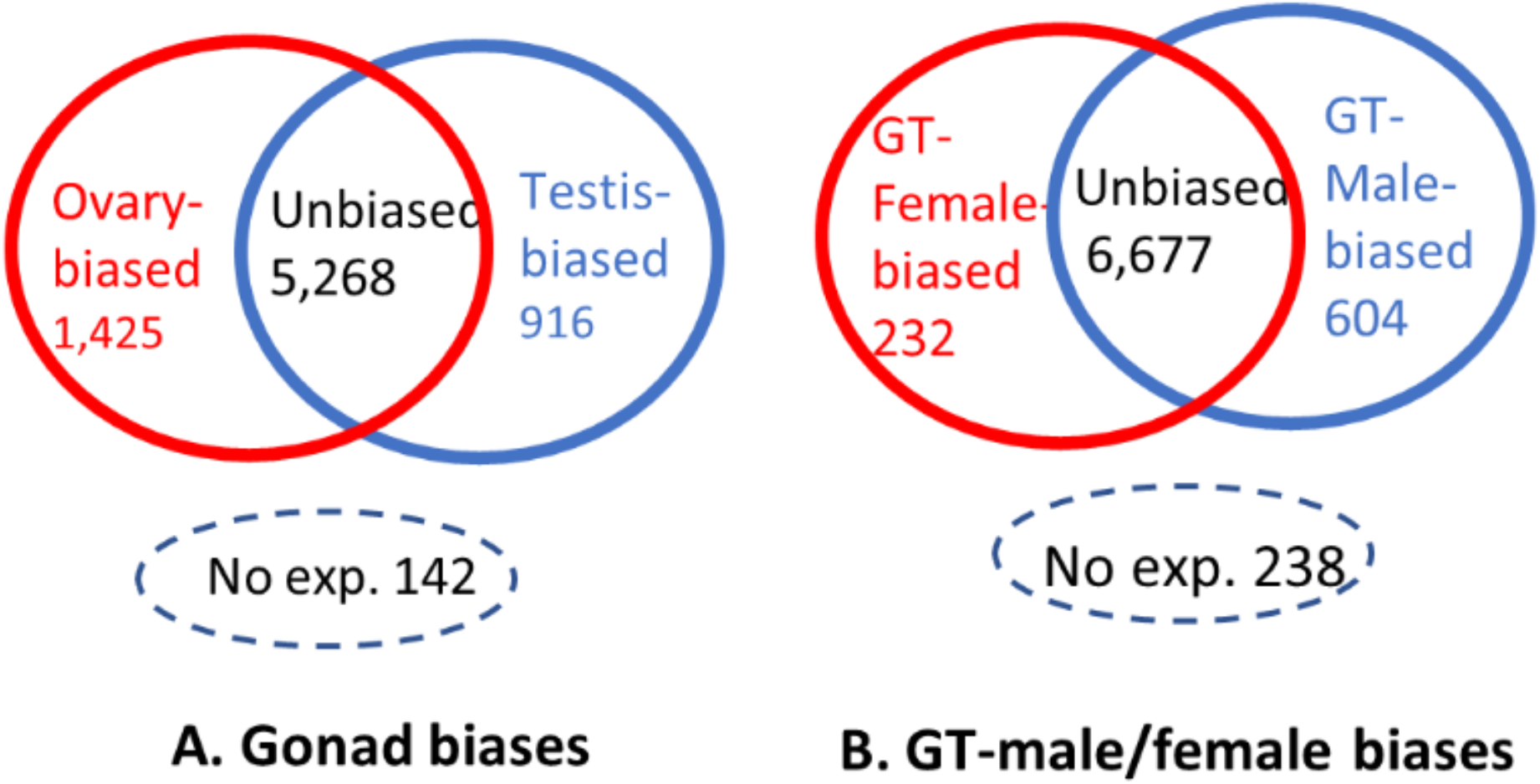
A Venn diagram showing the number of sex-biased and unbiased genes in the A) gonads; B) GT-soma.

**Fig. S3.**
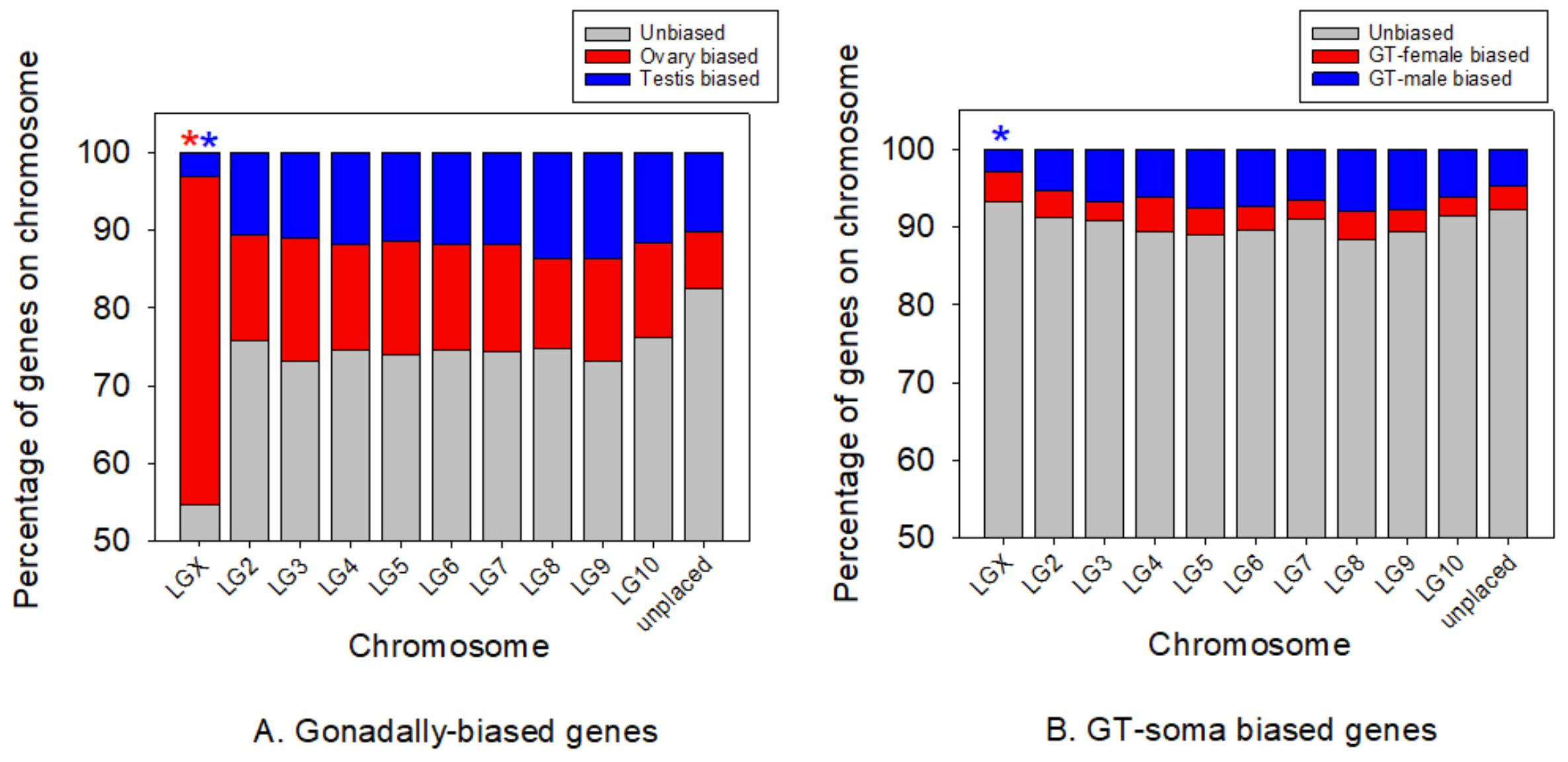
The frequency of sex-biased genes and unbiased genes on the X chromosome and autosome when all 16,434 genes of *T. castenum* are included in assessment. A) Gonadally biased genes; B) GT-soma biased genes. In A, the red and blue asterisks indicate more ovary-biased and fewer testis- (or fewer GT-male biased in B) biased genes respectively on the X-chromosomes than on pooled autosomes (Chi^2^-P with Yate’s correction P<0.05 for each contrast).

**Fig. S4.**
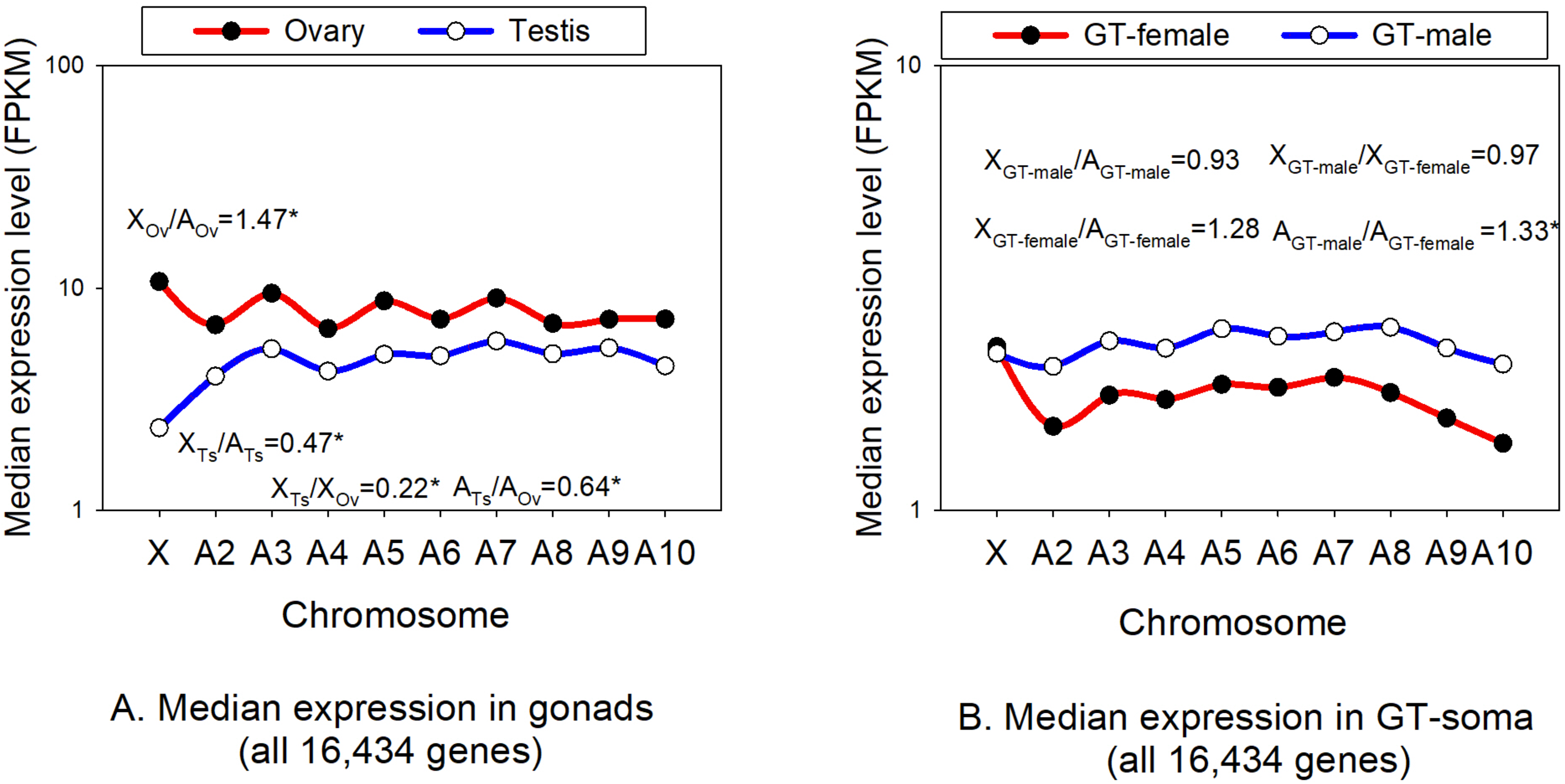
Median expression in the male and female tissues on each of the ten chromosomes in *T. castaneum* using all 16,434 genes in the genome. A) Gonads; B) GT-soma. For panel A, the ratio of median expression on the X chromosome (X) and autosomes (A) for testis-biased genes and for ovary-biased genes are shown (X_Ts_/A_Ts_ and X_Ov_/A_Ov_). Also shown are X_Ts_/X_Ov_ and A_Ts_/A_Ov_. Panel B contains the equivalent results for the GT-soma. Unmapped genes on chromosomes were excluded. *Indicates a statistically significant difference between the two groups contained in each ratio using MWU-tests.

**Table S1.** RNA-seq data used in present study before and after adapter and quality trimming with BBDuk (https://jgi.doe.gov/data-and-tools/bbtools/). The Short Read Archive (SRA) Biosample identifiers are also shown (https://www.ncbi.nlm.nih.gov/sra).

| Species Sample<sup>a</sup> | No. of Reads |  | SRA Biosample ID |
| --- | --- | --- | --- |
|  | Before trimming | After trimming |  |
| *Tribolium castaneum* |  |  |  |
| Testes sample 1 | 18,006,255 | 17,995,655 | SAMN12702873 |
| Ovary sample 1 | 39,140,493 | 39,122,050 | SAMN12702874 |
| GT-male sample 1 | 25,630,261 | 25,609,723 | SAMN12702875 |
| GT-female sample 1 | 41,513,717 | 41,472,348 | SAMN12702876 |
| Testes sample 2 | 24,795,583 | 24,787,238 | SAMN12702877 |
| Ovary sample 2 | 22,306,622 | 22,286,961 | SAMN12702878 |
| GT-male sample 2 | 62,781,001 | 62,712,242 | SAMN12702879 |
| GT-female 2 | 52,275,340 | 52,211,149 | SAMN12702880 |
| *Tribolium freemani* |  |  |  |
| Testes sample 1 | 32,222,092 | 32,203,312 | SAMN12702881 |
| Ovary sample 1 | 40,395,198 | 40,368,671 | SAMN12702882 |
| GT-male sample 1 | 32,933,858 | 32,926,509 | SAMN12702883 |
| GT-female sample 1 | 33,163,960 | 33,147,066 | SAMN12702884 |

<sup>a</sup> Reads were obtained from for two RNA-seq runs of each biological sample.

**Table S2.** **A.** The number of studied genes (N=7,751) with sex-biased expression in the gonads and GT-soma. B. The degree of overlap in sex-biased status on the X-linked genes between the gonads and GT-soma are also shown.

**A**
| N values gonads |  |  |  |  | N values GT-soma |  |  |  |  |
| --- | --- | --- | --- | --- | --- | --- | --- | --- | --- |
|  | Ovary-biased | Testis-biased | Unbiased<sup>a</sup> | Total |  | GT-female biased | GT-male biased | GT-unbiased<sup>a</sup> | Total |
| X-linked | 233 | 9 | 190 | 432 | X-linked | 24 | 12 | 396 | 432 |
| Autosomes | 1192 | 907 | 5220 | 7319 | Autosomes | 208 | 592 | 6519 | 7319 |
|  |  |  |  | 7751 |  |  |  |  | 7751 |

<sup>a</sup>Includes sexually unbiased genes and those with no expression in Fig. S2.

**B**
| Overlap on the X-Chromosome | N overlap in sex-biased status on the X-chromosome |
| --- | --- |
| Ovary-biased and GT-female biased | 17 |
| Testis-biased and GT-male biased | 0 |
| Ovary-biased and GT-unbiased | 213 |
| Testis-biased and GT-unbiased | 7 |
| GT-male biased and ovary-biased | 3 |

## Text File S1. Additional Methods

### Lab procedures

The gonads and other elements of the reproductive system were dissected as a single unit in ice cold 1x Phosphate Buffer Saline (PBS) and transferred immediately into TRIzol in a vial kept on dry ice. The reproductive tissues of males included the testes, accessory glands (mesadenia, ectadenia), vesicular seminalis, vas deferens and ejaculatory duct. The reproductive tissues for females included the ovaries, spermathecal gland, common oviduct, spermathecae, and vagina. All remaining nongonadal tissues of the adult body were collected and defined as GT-males and GT-females. Two biological samples per tissue type (testis, ovary, GT-males, GT-females) were collected for RNA-seq for our main target species for study, *T. castaneum* (eight total samples) while one sample per tissue type was obtained for *T. freemani* (four samples). A total of twelve samples were thus obtained for RNA-seq as shown in Table S1.

The testes, ovaries, GT-males and GT-females were stored in separate vials at −80°C until RNA extraction. RNA-isolation was performed according to the Ambion Life Technologies TRIzol Reagent Protocol, following which the RNA was used for RNA library preparation. Polyadenylated mRNAs were selected from total RNA samples using oligo-dT-conjugated magnetic beads on an Apollo324 automated workstation (PrepX PolyA mRNA isolation kit, Takara Bio USA). Entire poly-adenylated RNA samples were immediately converted into stranded Illumina sequencing libraries using 200 base pair (bp) fragmentation and sequential adapter addition on an Apollo324 automated workstation following manufacturer’s specifications (PrepX RNA-seq for Illumina Library kit, Takara Bio USA). Libraries were enriched and indexed using 14 cycles of amplification (LongAmp Taq 2x MasterMix, New England BioLabs Inc.) with PCR primers that included a 6bp index sequence to allow for multiplexing (custom oligo order from Integrated DNA Technologies). Excess PCR reagents were removed using magnetic bead-based cleanup on an Apollo324 automated workstation (PCR Clean DX beads, Aline Biosciences). Resulting libraries were assessed using a 2200 TapeStation (Agilent Technologies) and quantified by QPCR (Kapa Biosystems). Libraries were pooled and sequenced on two Illumina NextSeq 500 high output flow cells using single end, 75bp reads.

### Extracting CDS from *T. freemani*

To extract gene sequences from scaffolds in this species we used Web Augustus version 3.3.1 (http://bioinf.uni-greifswald.de/webaugustus/ [79]) that was trained to the *T. castaneum* genome and set at default parameters with the option to identify full length genes. The Augustus-generated CDS list for *T. freemani* was then assessed in ORF predictor, using its downloadable Perl script [80] to identify the highest quality reading frame per sequence. In ORF predictor, we employed the option to include the best-hit (lowest e-value) BLASTX alignment (conducted in BLAST+ v2.7.1, https://blast.ncbi.nlm.nih.gov) of *T. freemani* CDS versus the reference *T. castaneum* protein database to define reading frames, an approach which yielded 12,432 uninterrupted sequences of full or partial CDS for *T. freemani*. For further stringency in curating the *T. freemani* CDS list, we pooled the identified CDS with all 138,645,558 *T. freemani* RNA-seq reads (trimmed reads, Table S1) across all four tissue types (testis, ovaries, male carcass, female carcass) and mapped all sequences to the known and annotated CDS from *T. castaneum* using Geneious (v11.0.3), which generated consensus CDS. We then extracted CDS wherein all bases had a minimum of 10X coverage, and these were trimmed to the *T. castaneum* reference CDS. In those *T. freemani* CDS (obtained after ORF predictor) wherein the CDS was improved in quality (contained no unknown or ambiguous nucleotides) or in its length and/or the terminal stop codon was added by using the RNA-seq data, which occurred for N=1,249 CDS, we replaced the original Augustus-based CDS (among the 12,432) with the latter RNA-seq-mapped version CDS. Original CDS that were identified in *T. freemani* (and not found in the 12,432 list) only after using the RNA-seq mapping approach (N=196) were also included in the species final CDS list.

